# *B4GALT5*-deficient CHO-Lec2 cells expressing human α1,4-galactosyltransferase: a glycoengineered cell model for studying Shiga toxin receptors

**DOI:** 10.1101/2024.10.26.620404

**Authors:** Krzysztof Mikołajczyk

## Abstract

Human α1,4-galactosyltransferase (A4galt) is a glycosyltransferase existing in humans as two isoforms, widespread A4galt (named A4G) and its rare variant with p.Q211E substitution (A4Gmut). Both isoforms produce Gb3 (Galα1→4Galβ1→4Glc-Cer) on glycosphingolipids and P1 glycotope (Galα1→4Galβ1→4GlcNAc-R) on glycoproteins, which serve as receptors for Shiga toxin types 1 and 2 (Stx1 and Stx2). Stx1 is bound by Gb3 and P1 glycotope, while Stx2 is recognized solely by Gb3. To elucidate the role of these receptors, CHO-Lec2 cells expressing human A4G and A4Gmut were modified by disrupting the hamster *B4GALT5* gene using CRISPR/Cas9 technology. The *B4GALT5* gene encodes β1,4-galactosyltransferase 5 (B4galt5), synthesizing lactosylceramide, the key substrate for Gb3 synthesis. Consequently, *B4GALT5*-deficient CHO-Lec2-expressing A4G and A4Gmut cells lacked Gb3 glycosphingolipid but retained the ability to synthesize glycoprotein-based P1 glycotope. Both *B4GALT5*-deficient CHO-Lec2 cells expressing A4G and A4Gmut demonstrated no binding of Stx1B and Stx2B. The cytotoxicity assay showed that *B4GALT5*-deficient CHO-Lec2 cells expressing A4G were completely resistant to Stx1 holotoxin while A4Gmut-expressing cells revealed reduced sensitivity to Stx2. The glycoengineered CHO-Lec2 cells obtained in this study provide a valuable model for studying receptors for Stxs, enabling a detailed assessment of their roles in toxin binding and cytotoxicity.

## Introduction

Human α1,4-galactosyltransferase (A4galt, P1/P^k^ synthase, Gb3/CD77 synthase, EC 2.4.1.228) is encoded by the *A4GALT* gene. It plays a crucial role in the synthesis of Gb3 (P^k^, CD77, globotriaosylceramide, Galα1→4Galβ1→4Glc-Cer) and P1 (nLc5, Galα1→4Galβ1→4GlcNAcβ1→3Galβ1→4Glc-Cer), which are glycosphingolipids (GSLs) and, to a lesser extent, N-glycoproteins with terminal Galα1→4Galβ1→4GlcNAc-R structures (called P1 glycotopes) [1, 2, 3]. A4galt catalyzes the transfer of galactose from UDP-Gal to Gal-capped oligosaccharide acceptors. Two active isoforms of human α1,4-galactosyltransferase occur in humans. The most widespread A4galt isoform (A4G) synthesizes Gb3 and P1 on GSLs and P1 glycotope on GPs [4, 5, 6]. A rare enzyme variant (A4Gmut) contains a p.Q211E substitution (c.631C>G mutation in *A4GALT*, rs397514502) and can transfer Gal to GalNAc-terminating glycosphingolipids, forming NOR, NOR1 and NOR2 GSLs [2, 7]. The products of human A4galt belong to the P1PK histo-blood group system (International Society of Blood Transfusion ISBT No. 003) [7, 8, 9, 10, 11].

Gb3, the main A4galt enzyme product, is considered to be the main receptor for Shiga toxins (Stxs), released by *Shigella dysenteriae* serotype 1 and Shiga-producing strains of *Escherichia coli* (STEC). These strains secrete two distinct types of Shiga toxin, Stx1, which is homologous to *S. dysenteriae*-produced Stx and more genetically distinct Stx2. Both toxins are responsible for hemorrhagic colitis and life-threatening severe hemolytic-uremic syndrome (HUS), thus pose an increasing threat to human life [12, 13, 14]. Recently, it was found that glycoprotein-based P1 glycotope, alongside Gb3 glycosphingolipid, can bind Stx1 and mediate its cytotoxicity effects. Both receptors contain the Galα1→4Galβ structure, which is essential for the interaction with the B subunit of Stxs [4, 5]. Gb3 was also found to be associated with epithelial-to-mesenchymal transition, a process associated with cancerogenesis [15], while Gb3 accumulation is a hallmark of Fabry disease [16].

The dual acceptor specificity of human A4galt toward GSL- and GP-based acceptors may originate from its ability to form heterodimers with other GTs [17]. The primary substrate for human A4galt, LacCer is produced by LacCer synthase (β1,4-galactosyltransferase, B4galt), which occurs as two isoenzymes (β1,4-galactosyltransferase 5, B4galt5 and β1,4-galactosyltransferase 6, B4galt6) [18, 19]. Previous studies have shown the possibility of the existence of the A4galt-B4galt5 heterodimer by sharing by both enzymes the LXXR (X: any amino acid) sequence motifs [20, 21]. The presence of this motif in both A4galt and B4galt5 may indicate their close localization within the Golgi apparatus, facilitating heterodimer formation. However, the potential interactions between human A4galt and B4galt6 were suggested by Takematsu et al. (2011) [22]. Recently, using NanoBiT technology and computational analysis of Alphafold enzyme models, it was demonstrated that A4galt can form homodimers and heterodimers with human B4galt1 and B4galt5 [17].

In a previous study, CHO-Lec2 cells were transfected with the human *A4GALT* gene, leading to the synthesis of Gb3 and P1 glycotope. The use of CHO-Lec2 cells was motivated by the lack of a CMP-sialic acid transporter, which is essential for sialic acid-containing glycan synthesis [23, 24]. This feature prevents interference from sialylated glycans in Stx binding to cell surface receptors [1, 5]. Using CHO-Lec2 cells, it was found that glycotopes containing the P1 epitope (Galα1→4Galβ1→4GlcNAc-R) can be functional receptors for Stx1, in addition to the canonical GSL-based Gb3 receptor [4, 5]. Herein, we eliminated the Gb3 GSL receptor by disrupting the hamster *B4GALT5* gene using CRISPR/Cas9 [25]. *B4GALT5* encodes hamster B4galt5, a major enzyme isoform (alongside B4galt6), responsible for the galactosylation of glycans in CHO-Lec2 cells. Furthermore, considering the ability of human A4galt to recognize two different acceptors and the A4galt-B4galt5 heterodimer formation in CHO-Lec2 cells [17], it was aimed to generate *B4GALT5*-deficient CHO-Lec2 A4G and A4Gmut cells. Therefore, these glycoengineered cells were analyzed to assess Stx binding and sensitivity to Stx1- and Stx2-mediated cytotoxicity.

## Material and methods Antibodies

Human anti-P1 (clone P3NIL100, recognizing only P1 antigen and P1 glycotope), mouse anti-P1 (clone 650, recognizing Gb3 and P1 antigens), anti-6x-His Tag antibody (clone HIS.H8), and biotinylated goat anti-mouse IgG/A/M (H/L) antibodies were purchased as described in [5]. Goat anti-human polyvalent immunoglobulin biotinylated antibody and alkaline phosphatase-ExtrAvidin were obtained from Sigma-Aldrich, St. Louis, MO. The recombinant His-tagged Stx1B and Stx2B subunits were used in western blotting and flow cytometry analysis, as described in [5]. Stx1 (Verotoxin 1, from *E. coli* O157) and Stx2 (Verotoxin 2, from *E. coli* O157) holotoxins for cytotoxicity assay were purchased from Sigma-Aldrich, St. Louis, MO, USA.

### Generation *B4GALT5*-deficient CHO-Lec2 cells-expressing human A4galt using CRISPR/Cas9

CHO-Lec2 cells were transduced with *A4GALT* and *A4GALT* with c.631C>G ORFs genes, encoding A4galt (designated as A4G) and A4galt with p.Q211E substitution (A4Gmut), respectively, as described previously [5]. For specific experiments, CHO-Lec2 A4G, CHO-Lec2 A4Gmut, and untransfected cells were cultured for 2 weeks in a complete medium containing 3 μM Genz-123346 (Merck, Darmstadt, Germany), a glucosylceramide synthase inhibitor [26].

Guide RNA sequences for the CRISPR/Cas9 system were designed as described in [27] with the use of CRISPOR software (http://crispor.tefor.net/crispor.py, [28]). All obtained sgRNAs were cloned into pSpCas9(BB)-2A-GFP (PX458, a gift from Feng Zhang, Addgene plasmid #48138; http://n2t.net/addgene:48138; RRID:Addgene_48138, Addgene plasmid #48138) [26]. The ligated products were transformed into electrocompetent *E. coli* DH5α cells (Thermo Fisher Scientific, Waltham, MA, USA), and the cells were plated onto ampicillin selection plates (100 μg/mL ampicillin) and incubated at 37°C overnight. Plasmid DNA was isolated using the Gene JET Plasmid Miniprep kit (Thermo Fisher Scientific, Waltham, MA, USA) and verified by Sanger sequencing. Nucleotide sequences of sgRNA are listed in Table SIA.

### Sanger sequencing of mutated sites

*B4GALT5* gene disruption was assessed using a PCR-based method (Laboratory of DNA sequencing and synthesis, Institute of Biochemistry and Biophysics PAS, Warsaw). Briefly, purified PCR products of *B4GALT5* were cloned into pCRScript Amp SK (Promega, Madison, WI) and transformed into *E. coli* Top10 competent cells. Using the positive colonies as a template, a second PCR reaction with M13Forward and M13Reverse standard primers was performed (Table SIB). The obtained nucleotide fragments of *B4GALT5* were purified and sequenced. Mutations were identified by alignment of the sequenced alleles with the wild-type gene reference NW_003613899.1 (CriGri_1.0, scaffold2107) using GeneStudio Professional software.

### Western blotting

CHO-Lec2 cells were lysed in CelLytic M (Thermo Fisher Scientific, Waltham, MA, USA) supplemented with a protease inhibitor cocktail. In total, 20 μg of cell lysate per lane was subjected to nonreducing sodium dodecyl sulfate-polyacrylamide gel electrophoresis (SDS-PAGE) on a 10% Tris-Glycine gel and transferred to a nitrocellulose membrane as described in [1].

### HPTLC orcinol staining

Orcinol staining was performed using standard procedures as previously described [1]. Briefly, dried thin-layer chromatography (HPTLC) plates were sprayed with an orcinol solution (0.2% w/v) in 3 M aqueous sulfuric acid and incubated in an oven at 110°C for 10 min.

### Extraction and purification of glycosphingolipids from CHO-Lec2 cells

Isolation, fractionation, and HPTLC analysis of CHO-Lec2 neutral GSLs were performed as described [1]. Briefly, GSLs were extracted with chloroform/methanol, desalted and separated from phospholipids and gangliosides. GSLs solubilized in chloroform/methanol (2:1, v/v) were applied to HPTLC plates (Kieselgel 60, Merck, Darmstadt, Germany) and developed in chloroform/methanol/water (55:45:9, v/v/v). Dried plates were stained with orcinol or prepared for antibody assay according to [1].

### MALDI-TOF mass spectrometry analysis of GSLs extracted from CHO-Lec2 cells

Matrix-assisted laser desorption/ionization-time of flight (MALDI-TOF) mass spectrometry was carried out on an ultrafleXtreme MALDI TOF/TOF instrument (BrukerDaltonics, Bremen, Germany), with the use of Peptide Calibration Standard (BrukerDaltonics, Bremen, Germany) as external calibration and Norharmane (9HPyrido[3,4-b]indole, Sigma-Aldrich, St. Louis, MO) as a matrix (10 mg/ml, chloroform:methanol, 2:1, v/v). GSLs were dissolved in chloroform/ methanol (2:1, v/v). Spectra were scanned in the range of *m/z* 700–2000 in the reflectron-positive mode.

### Quantitative analysis of *A4GALT* transcript in CHO-Lec2 cells

Total RNA from transfected or non-transfected CHO-Lec2 cells was prepared using Universal RNA Purification Kit (Eurx, Gdansk, Poland) and the complementary DNAs (cDNAs) were synthesized using the SuperScript III First-Strand Synthesis kit (Life Technologies, Carlsbad, CA, USA) with oligo(dT) primers. A quantitative polymerase chain reaction (qPCR) was performed with 30 ng of cDNA using the CFX Connect qPCR system (Bio-Rad, Hercules, CA, USA), according to the manufacturer’s instruction. Human *A4GALT* and hamster *B4GALT5* transcripts were detected using the Custom TaqMan Gene Expression Assay. For endogenous control normalization, the custom assay targeting *C. griseus* (Chinese hamster) *GAPDH* was used. All samples were run in triplicate. The target nucleotide sequences are shown in Table SIC, and the RT-qPCR conditions are presented in Table SII.

### Flow cytometry

CHO-Lec2 cells were scraped, washed (with PBS), and incubated for 30 min on ice with anti-P1 (clone P3NIL100; 1:400; all dilutions were done with 1% BSA in PBS), anti-P1 (clone 650; 1:200) antibodies, washed, and incubated with secondary FITC-conjugated antibodies: anti-human IgM (1:100), anti-mouse IgM (1:100). To analyze Shiga toxin binding, cells were incubated with 1 μg/ml Stx1B or Stx2B, then washed, and incubated with anti-6x-His Tag antibody (1:1000), followed by washing and incubation with FITC-conjugated anti-mouse IgG (1:100). After incubation with FITC-conjugates, the cells were washed, resuspended in 500 μl of cold PBS, and subjected to flow cytometry analysis using FACSCalibur (BD Biosciences, Franklin Lakes, NJ, USA). In total, 1 × 10^5^ events from the gated population were analyzed by Flowing Software (Perttu Terho, University of Turku). The antibodies and StxB subunits binding capacity per cell of CHO-Lec2 was determined using Quantum (Bio-Rad, Hercules, CA, USA) as described in [1]. For statistical analysis, one-way ANOVA with Bonferroni post-hoc test was used. All analyses were performed with GraphPad Prism (GraphPad Software, CA, USA). Statistical significance was assigned to *p*-value <0.05.

### Cytotoxicity assay

In total, 2 × 10^4^ CHO-Lec2 cells were seeded in 96-well plates (Wuxi NEST Biotechnology Co., Ltd, China) in complete DMEM/F12. After 24 h medium was replaced with 100 μl/well of serum-free DMEM/F12 containing 0.01, 0.05, 0.1, 0.5, and 1 ng/ml of Stx1 or Stx2 holotoxins (all concentrations were run in triplicate). After 20 h of toxin treatment, 20 μl/well of MTS tetrazolium compound (CellTiter 96 AQueous One Solution Assay, Promega, Madison, WI) was added as described in [1]. The background absorbance registered at zero cells/well was subtracted from the data and the absorbance of wells incubated in medium without the addition of Stx was taken as 100% of cell viability.

## Results

### Generation CHO-Lec2 A4galt-expressed cells with disrupted *B4GALT5* gene

Genome sequence analysis of CHO-Lec2 A4G and A4Gmut with *B4GALT5* KO showed heterozygotic complex mutations within exon 5 of the *B4GALT5* gene (Fig. S1). The mutations identified included one deletion and two insertions of nucleotides within *C. griseus B4GALT5* in CHO-Lec2 A4G cells (Fig. S1A), and one deletion and one insertion in CHO-Lec2 A4Gmut cells (Fig. S1C). These nucleotide aberrations led to significantly reduced *B4GALT5* gene transcript in CHO-Lec2 A4G and A4Gmut with *B4GALT5* KO (Fig. S1B, D). These findings indicated the effective disruption of the *B4GALT5* gene in CHO-Lec2 expressing both A4G and A4Gmut.

### Shiga toxins binding to CHO-Lec2 cells expressing A4G and A4Gmut with *B4GALT5* KO

Shiga toxin types 1 and 2 are known to bind to the Galα1→4Gal epitope, located on the Gb3 GSL and GP-based P1 glycotope on the cell surface. It was anticipated that the generated CHO-Lec2 cells expressing A4G and A4Gmut with *B4GALT5* KO lacked globo-series GSL synthesis, including Gb3. To further confirm the depletion of Gb3 in these modified cells, the CHO-Lec2 A4G and A4Gmut cells (*B4GALT5* KO and with active gene) were treated with Genz-123346 [5].

To evaluate the presence of Gb3 GSLs, the glycolipids isolated from CHO-Lec2 A4G and A4Gmut cells (with or without *B4GALT5* KO) were analyzed by HPTLC and MALDI-TOF mass spectrometry. Orcinol staining (which recognizes glycolipids) exhibited no neutral GSLs in CHO-Lec2 A4G and A4Gmut *B4GALT5 KO* cells, similar to the cells treated with Genz-123346 (Fig. 1). Analysis of GSLs using HPTLC overlaid with anti-P1 (clone 650) antibody showed no bands corresponding to Gb3Cer and Gb4Cer GSLs in CHO-Lec2 A4G and A4Gmut *B4GALT5* KO cells (Fig. 1). Using MALDI-TOF mass spectrometry, only HexCer isoforms were found in both CHO-Lec2 A4G and A4Gmut *B4GALT5* KO cells (Fig. S2), which corresponded to the HPTLC overlay and orcinol staining results. Taken together, CHO-Lec2 A4G and A4Gmut *B4GALT5* KO cells were Gb3 GSL-deficient.

**Fig. 1.**
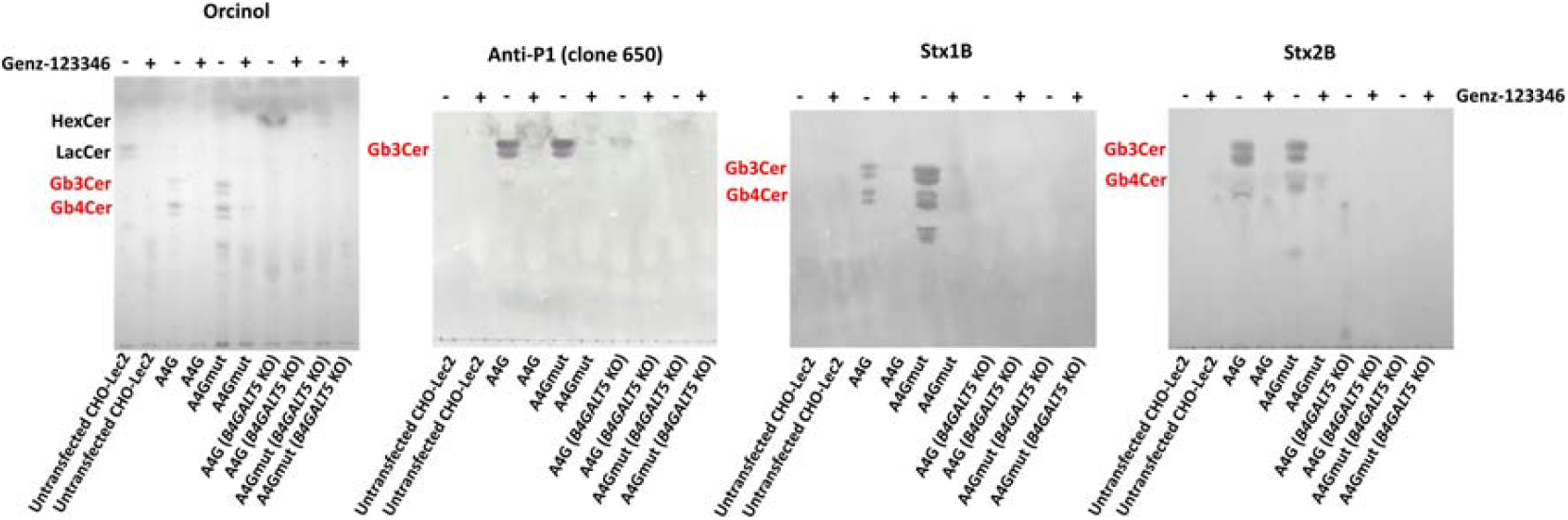
HPTLC analysis of neutral glycosphingolipids purified from CHO-Lec2 cells expressing human A4G and A4Gmut, with or without hamster *B4GALT5* KO. The GSL samples were stained with orcinol, anti-P1 (clone 650), Stx1B and Stx2B. CHO-Lec2 A4G and A4Gmut cells with active *B4GALT5* gene produced GSLs, including HexCer, LacCer, Gb3Cer and Gb4Cer. Among them, Gb3Cer and Gb4Cer (marked in red) were bound by Stx1B and Stx2B. A4G, CHO-Lec2 cells expressing human A4galt; A4Gmut, CHO-Lec2 cells expressing human A4galt with p.Q211E; *B4GALT5* KO, CHO-Lec2 A4G/A4Gmut with hamster *B4GALT5* knock-out.

Quantitative flow cytometry was performed to evaluate the binding capacity of anti-P1 (clones P3NIL100 and 650) and the Stx1B and Stx2B to CHO-Lec2 A4G and A4Gmut cells. CHO-Lec2 A4G and A4Gmut cells with *B4GALT5* KO did not show Stx1B or Stx2B binding to the cell surface (Fig. 2). Treatment of CHO-Lec2 A4G and A4Gmut *B4GALT5* KO cells with the Genz-123346 inhibitor also resulted in no Stx1B and Stx2B binding (Fig. 2). Despite abolished Stxs binding to the cells, CHO-Lec2 A4G and A4Gmut *B4GALT5* KO cells still bound anti-P1 antibodies (cloned 650 and P3NIL100), which recognize molecules containing the Galα1→4Gal epitope (Fig. 2). In summary, CHO-Lec2 A4G and A4Gmut *B4GALT5* KO cells displayed no binding of Stx1B and Stx2B to the cell surface, along with the retention of molecules bound by anti-P1 antibodies.

**Fig. 2.**
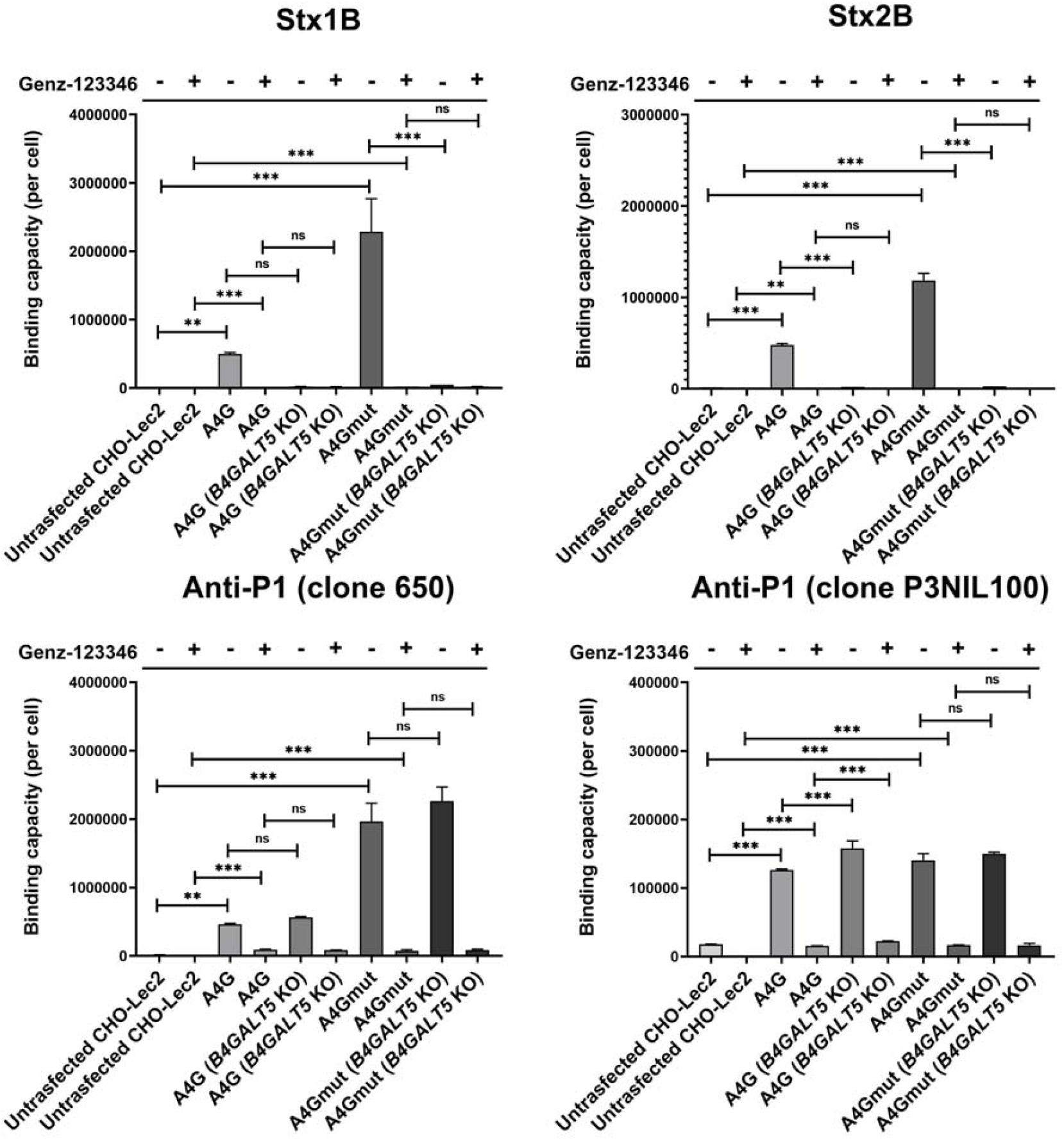
Quantitative flow cytometry analysis of the anti-P1 (clones 650 and P3NIL100), Stx1B and Stx2B binding capacity of CHO-Lec2 A4G and A4Gmut cells. The cells were either untreated or treated with Genz-123346 inhibitor. The data are presented as medians from at least three replicates with error bars representing interquartile ranges. Statistical significance was established as *, p < 0.05; **p < 0.01; ***p < 0.001; ns, not significant. A4G, CHO-Lec2 cells expressing human A4galt; A4Gmut, CHO-Lec2 cells expressing human A4galt with p.Q211E; *B4GALT5* KO, CHO-Lec2 A4G/A4Gmut with hamster *B4GALT5* knock-out.

Western blotting was conducted to evaluate GP-based P1 glycotope Stx receptor in CHO-Lec2 A4G and A4Gmut (with and without *B4GALT5* KO) cell lysates using the anti-P1 antibody (clone P3NIL100) and the Stx1B and Stx2B. No significant differences were observed between the analyzed cells (regardless of the presence *B4GALT5* KO or not) (Fig. 3). However, CHO-Lec2 A4Gmut cells exhibited higher levels of P1-based glycoproteins, consistent with previous findings [5]. Additionally, the binding of Stx B subunits to cell lysates showed that Stx1B binds to several glycoproteins, while Stx2B did not detect any proteins, regardless of *B4GALT5* deficiency (Fig. 3). Treatment with Genz-123346 did not significantly affect the binding patterns of the anti-P1 antibody (clone P3NIL100) and Stx1B to CHO-Lec2 A4G and A4Gmut (*B4GALT5* KO and with the active gene) cell lysates (Fig. 3). Collectively, CHO-Lec2 A4G and A4Gmut (irrespective of *B4GALT5* deficiency) still produced P1 glycotope on GPs, which are bound by Stx1B.

**Fig. 3.**
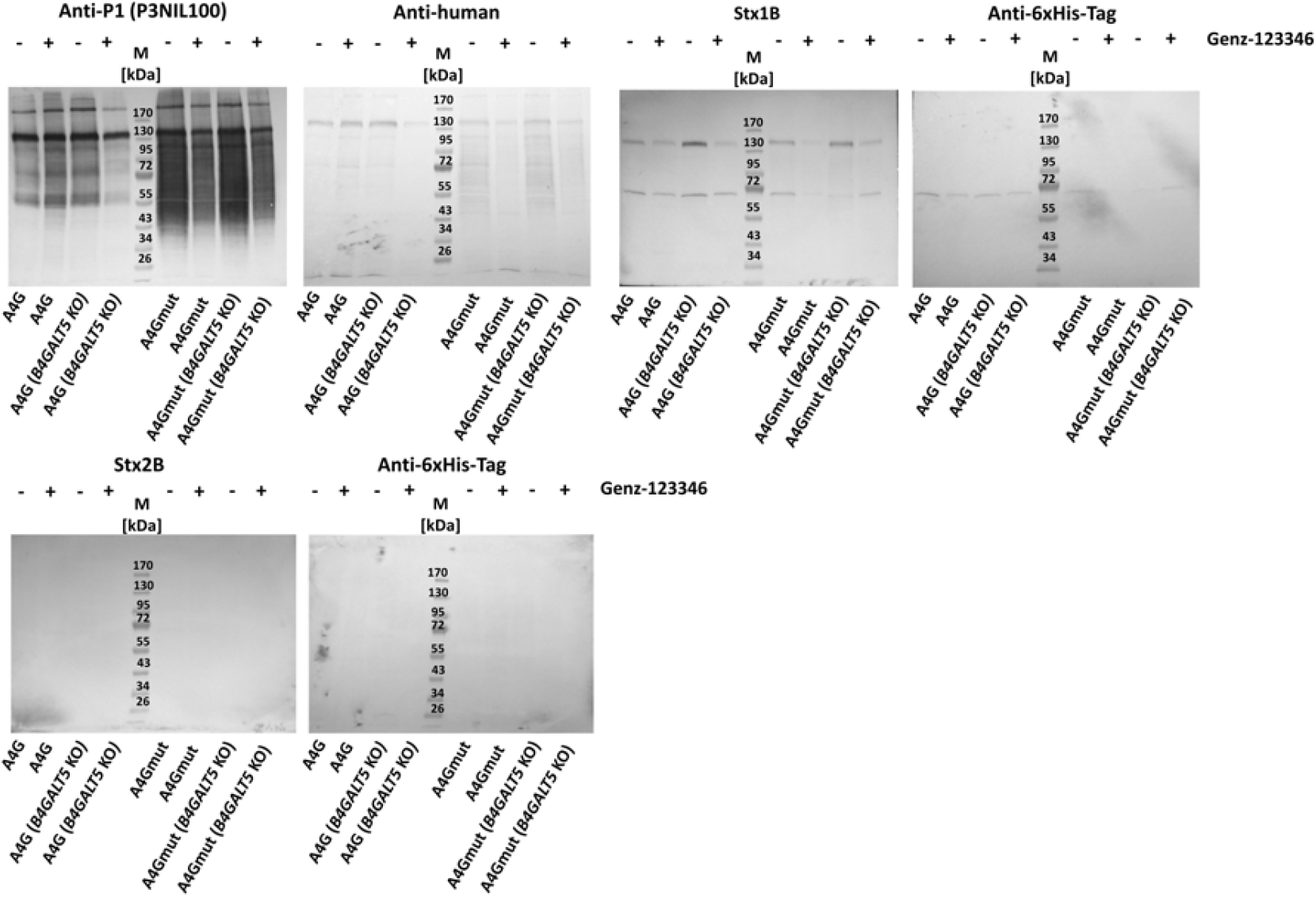
Western blotting analysis of cell lysates from CHO-Lec2 expressing human A4G and A4Gmut, with or without hamster *B4GALT5* KO. The cells were cultured with or without Genz-123346 inhibitor and stained with an anti-P1 antibody (clone P3NIL100). A4G, CHO-Lec2 cells expressing human A4galt; A4Gmut, CHO-Lec2 cells expressing human A4galt with p.Q211E; *B4GALT5* KO, CHO-Lec2 A4G/A4Gmut with hamster *B4GALT5* knock-out.

### Sensitivity of CHO-Lec2 cells expressing A4G and A4Gmut with *B4GALT5* KO for Shiga holotoxins

To assess how *B4GALT5* disruption affects the viability of CHO-Lec2 A4G and A4Gmut cells upon Stx holotoxin exposure, a cytotoxicity assay was performed. Untransfected CHO-Lec2 cells, which are resistant to both Stx1 and Stx2, were used as controls. *B4GALT5*-deficient CHO-Lec2 A4G cells were not sensitive to either Stx1 or Stx2 treatment, while CHO-Lec2 A4Gmut *B4GALT5* KO cells were resistant to Stx1 but slightly sensitive to Stx2 (Fig. 4A). When treated with the Genz-123346 inhibitor, both CHO-Lec2 A4G and A4Gmut *B4GALT5* KO cells showed no cytotoxic effects in response to either Stx1 or Stx2 (Fig. 4B). In summary, *B4GALT5*-deficient CHO-Lec2 A4G cells were resistant to both holotoxins, whereas CHO-Lec2 A4Gmut *B4GALT5* KO cells demonstrated a slight sensitivity to Stx2 but remained unaffected by Stx1. These results suggest the presence of an unidentified receptor for Stx2 holotoxin (probably belonging to the GSLs), produced exclusively in CHO-Lec2 A4Gmut cells.

**Fig. 4.**
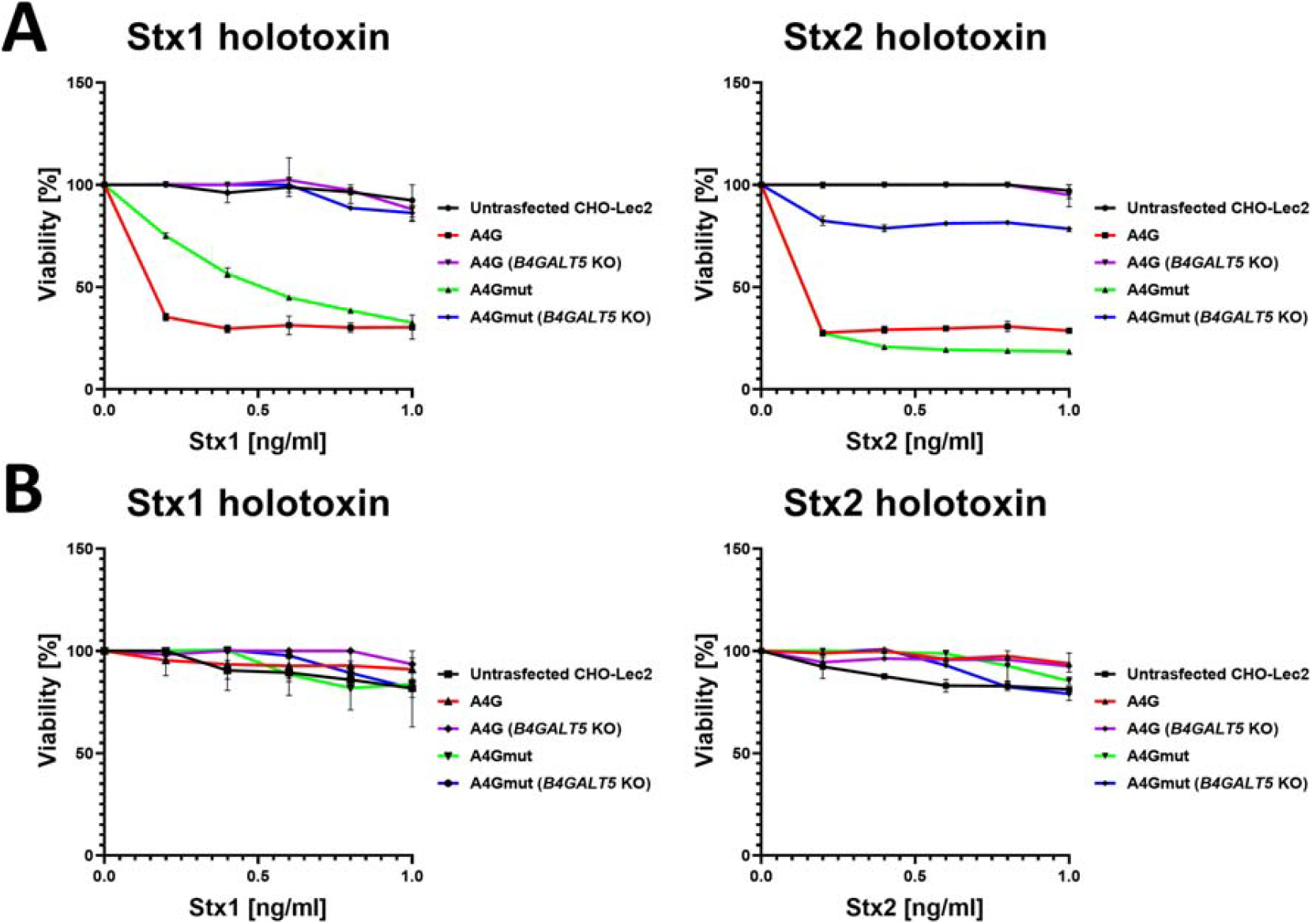
Shiga toxins cytotoxicity assay. The viability of CHO-Lec2 cells expressing human A4G and A4Gmut, with or without hamster *B4GALT5* KO, was analyzed after exposure to Stx1 or Stx2 in the absence (A) or presence (B) of Genz-123346 inhibitor. A4G, CHO-Lec2 cells expressing human A4galt; A4Gmut, CHO-Lec2 cells expressing human A4galt with p.Q211E; *B4GALT5* KO, CHO-Lec2 A4G/A4Gmut with hamster *B4GALT5* knock-out.

## Discussion

In this study, we obtained *B4GALT5*-deficient CHO-Lec2 A4G and A4Gmut-expressing cells lacking Gb3 GSLs which provides a good model for investigating glycoprotein-based Stx receptors. In recent years, the development of glycoengineered mammalian cells using chemical, enzymatic, and genetic approaches, including CRISPR/Cas9-based gene editing, has become increasingly important in biomedicine [25]. Gene silencing, especially these encoding GTs, has been widely used to modify cell surface receptors involved in pathogen recognition and modulation of the immune response. For example, silencing *ST6GAL1* expression in airway epithelial cells using siRNA, reduced the binding of human influenza viruses to cell surface sialic acid-containing receptors [29].

Cells with modified glycosylation machinery provide insights into glycan functions and molecular interactions and serve as valuable tools for designing cells with specific glycans on their surface [30, 31]. The *B4GALT5*-deficient CHO-Lec2 A4G and A4Gmut cells obtained in this study could be utilized for further research on the dual acceptor specificity of human A4galt. For years, Gb3 has been considered the primary receptor for Stxs, however, recent findings indicated that Stx receptors are located on GSLs and GPs, with different binding patterns for Stx1 and Stx2 [5]. Studies on pigeon A4galt homologs have revealed the existence of two enzymes with distinct acceptor specificities: one primarily utilizes GPs and the other recognizes both GSLs and GPs as substrates [6]. Similarly, human A4galt has demonstrated dual specificity toward GSLs (major substrates) and GPs (minor substrates) [5]. This phenomenon, referred to as acceptor promiscuity, has been well documented within GTs and has significant implications for biological and pathological processes [32]. Examples of such promiscuity include human A and B transferases which synthesize A and B blood group antigens on both N- and O-glycans as well as GSLs [33]. Similarly, human α1,3-fucosyltransferases (Futs) demonstrate isoenzyme-dependent specificity, with certain isoforms targeting glycolipids and others preferentially glycoprotein acceptors [34].

The analysis of *B4GALT5*-deficient CHO-Lec2 cells confirmed that Gb3 is the primary functional receptor for Stx1 and Stx2. Notably, due to the absence of Gb3 GSLs, this cell line may serve as a good model for investigating N-glycoprotein Stx receptors. Previous studies have explored the relationship between Stxs resistance and GTs knock-out (including *A4GALT* and *B4GALT5*) in various human cells, such as HeLa [35, 36], bladder cancer cells 5637 [37] and colorectal adenocarcinoma cells HT-29 cells [38]. For example, Tian et al. 2018 showed that the bladder carcinoma cell line 5637 cells with *B4GALT5* knock-out revealed increased resistance to both Stx1 and Stx2, compared to the control cells [37]. More comprehensive studies are needed to investigate the novel receptor for Stx2 holotoxin, which was identified in *B4GALT5*-deficient CHO-Lec2 A4Gmut cells (Fig. 4).

*B4GALT5*-deficient CHO-Lec2 cells could be further glycoengineered by modulating the synthesis of GP-based receptors or altering the cell surface glycome by the introduction of genes encoding specific GTs. Future research should focus on elucidating the dual specificity of A4galt toward GSLs and GPs and understanding the molecular mechanisms governing the distribution of Stx receptors on different acceptor molecules.

## Abbreviations

A4galt: α1,4-galactosyltransferase, Gb3/CD77 synthase
B4galt: β1,4-galactosyltransferase
Gb3: globotriaosylceramide, P^k^ antigen, CD77, Galα1→4Galβ1→4Glc-Cer
Genz-123346: *N*-[(1*R*,2*R*)-1-(2,3-dihydro-1,4-benzodioxin-6-yl)-1-hydroxy-3pyrrolidin-1-ylpropan-2-yl]nonanamide, glucosylceramide synthase inhibitor
GP: glycoprotein
GSL: glycosphingolipid
HPTLC: thin-layer chromatography
HUS: hemolytic-uremic syndrome
MALDI-TOF: matrix-assisted laser desorption/ionization-time of flight
LacCer: lactosylceramide
P1: nLc5, Galα1→4Galβ1→4GlcNAcβ1→3Galβ1→4Glc-Cer
P1 glycotope: terminal Galα1→4Galβ1→4GlcNAc-R disaccharide on N-glycoproteins
STEC: Shiga toxin-producing *Escherichia coli*
Stx: Shiga toxin
Stx1B: B subunit of Shiga toxin 1
Stx2B: B subunit of Shiga toxin 2

## Acknowledgments

This research was funded by the National Science Centre of Poland, PRELUDIUM 20 Project 2021/41/N/NZ6/00949. The graphical abstract was created with BioRender.com.

