## Supplementary Material for "*B4GALT5*-deficient CHO-Lec2 cells expressing human α1,4-galactosyltransferase: a glycoengineered cell model for studying Shiga toxin receptors"

Krzysztof Mikołajczyk^1,2^

^1^ Laboratory of Glycobiology, Hirszfeld Institute of Immunology and Experimental Therapy, Polish Academy of Sciences, Rudolfa Weigla St. 12, 53-114, Wroclaw, Poland

^2^ Correspondence author:

Krzysztof Mikołajczyk

Rudolfa Weigla St. 12, 53-114 Wroclaw, Poland

**Contents**

**Fig. S1**. Generation of *B4GALT5*-deficient CHO-Lec2 cells expressing human A4galt (A4G, A and B) and A4galt with p.Q211E (A4Gmut, C and D). Sanger sequencing was used to verify the knock-out of hamster *B4GALT5* (A and C). RT-qPCR was used to evaluate the hamster *B4GALT5* transcript levels in the CHO-Lec2 cells with and without *B4GALT5* KO (B and D).

**Fig. S2.** MALDI-TOF analysis of neutral glycosphingolipids isolated from CHO-Lec2 cells expressing human A4galt (A4G) and A4galt with p.Q211E (A4Gmut), cultured in the presence or absence of the Genz-123346 inhibitor. CHO-Lec2 A4G and A4Gmut cells, with active *B4GALT5* gene, untreated with Genz-123346 produced hexosylceramide (HexCer), lactosylceramide (LacCer), globotriaosylceramide (Gb3Cer) and globotetraosylceramide (Gb4Cer), while cells treated with Genz-123346 synthesized only HexCer (probably galactosylceramide, due to inhibition of GlcCer synthesis by Genz-123346). For CHO-Lec2 A4G and A4Gmut cells with *B4GALT5* KO, only HexCer was detected, regardless of the presence or absence of Genz-123346 in the culture medium. *B4GALT5* WT, CHO-Lec2 A4G/A4Gmut cells without *B4GALT5* gene knock-out; *B4GALT5* KO, CHO-Lec2 A4G/A4Gmut cells with *B4GALT5* gene knock-out.

**Table SI. A)** Nucleotide sequences used for generation of sgRNAs to knock-out hamster *B4GALT5* gene in CHO-Lec2 cells. **B)** Nucleotide sequences of primers used in Sanger sequencing.

**Table SII.** RT-qPCR conditions used for quantitative analysis of *A4GALT* and *B4GALT5* transcripts.

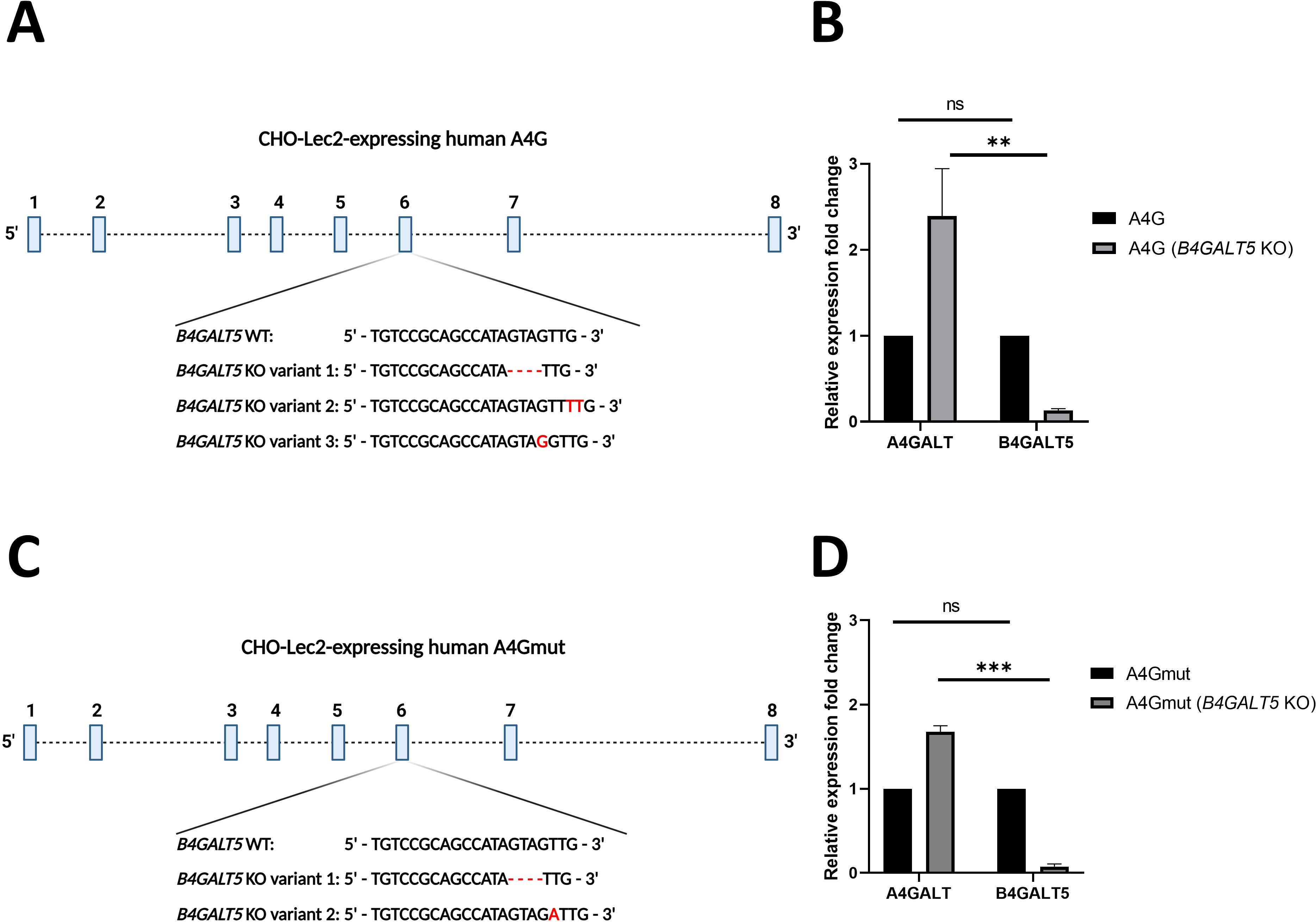

**Fig. S1.** Generation of *B4GALT5*-deficient CHO-Lec2 cells expressing human A4galt (A4G, A and B) and A4galt with p.Q211E (A4Gmut, C and D). Sanger sequencing was used to verify the knock-out of hamster *B4GALT5* (A and C). Real-time qPCR was used to evaluate the hamster *B4GALT5* transcript levels in the CHO-Lec2 cells with and without *B4GALT5* KO (B and D).

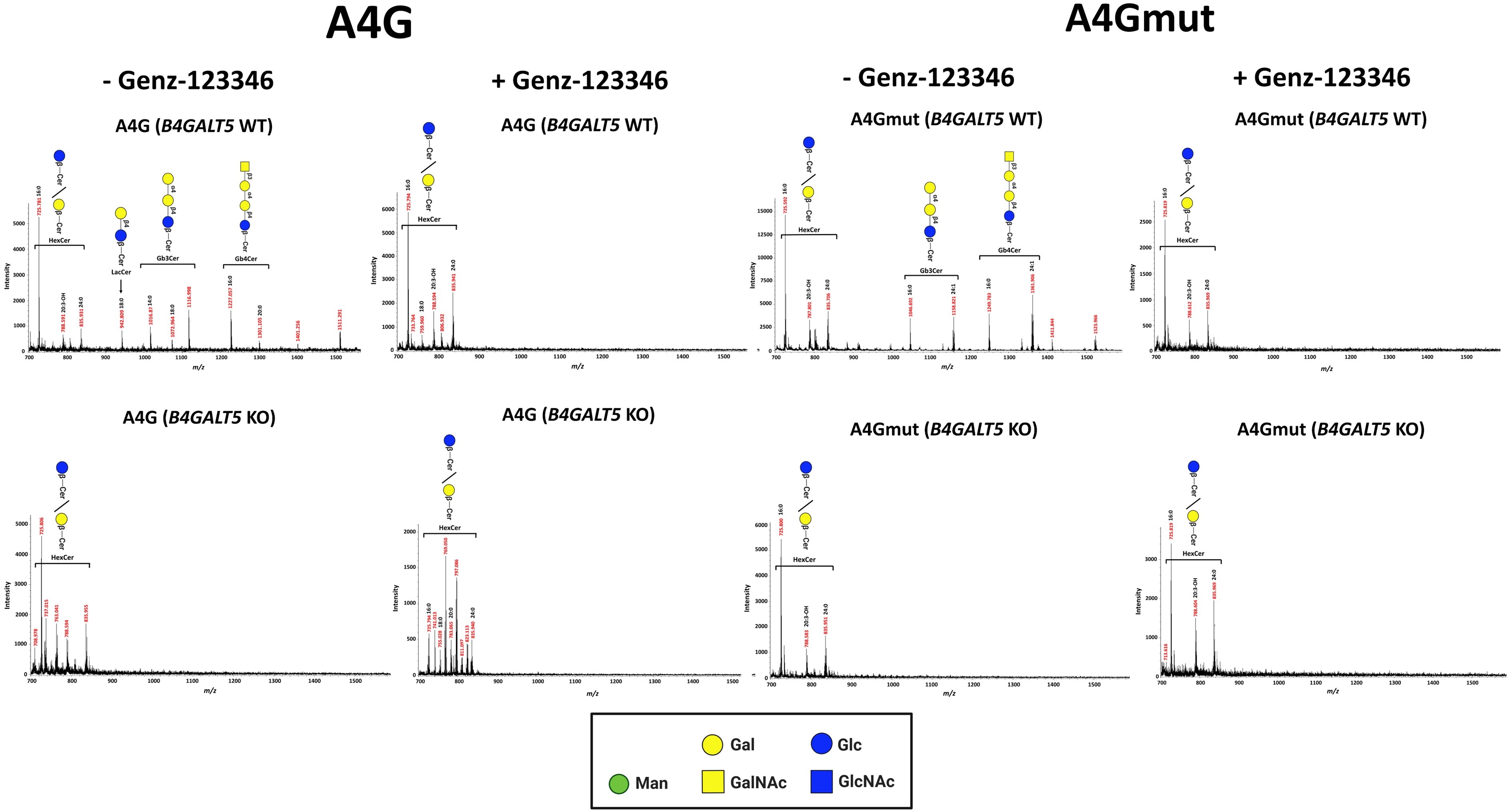

**Fig. S2.** MALDI-TOF analysis of neutral glycosphingolipids isolated from CHO-Lec2 cells expressing human A4galt (A4G) and A4galt with p.Q211E (A4Gmut), cultured in the presence or absence of the Genz-123346 inhibitor. CHO-Lec2 A4G and A4Gmut cells, with active *B4GALT5* gene, untreated with Genz-123346 produced hexosylceramide (HexCer), lactosylceramide (LacCer), globotriaosylceramide (Gb3Cer) and globotetraosylceramide (Gb4Cer), while cells treated with Genz-123346 synthesized only HexCer (probably galactosylceramide GalCer, due to inhibition of glucosylceramide GlcCer synthesis by Genz-123346). For CHO-Lec2 A4G and A4Gmut cells with *B4GALT5* KO, only HexCer was detected, regardless of the presence or absence of Genz-123346 in the culture medium. *B4GALT5* WT, CHO-Lec2 A4G/A4Gmut cells without *B4GALT5* gene knock-out; *B4GALT5* KO, CHO-Lec2 A4G/A4Gmut cells with *B4GALT5* gene knock-out.

**Table SI.** Oligonucleotides used in this study.

**A)** Nucleotide sequences used for generation of sgRNAs to knock-out hamster *B4GALT5* gene in CHO-Lec2 cells.

| **sgRNA** | **Nucleotide sequence (5’ – 3’)** |
| --- | --- |
| sgRNA_Exon5_s | CACCGTCCGCAGCCATAGTAGTTG |
| sgRNA_Exon5_a | AAACCAACTACTATGGCTGCGGAC |

underlined, additional nucleotide overhangs according to Ran et al. (2013) [26].

**B) Nucleotide sequences of primers used in Sanger sequencing.**

| **Name** | **Nucleotide sequence (5’ – 3’)** |
| --- | --- |
| **Sequencing primers** | |
| M13_forward | GTAAAACGACGGCCAGT |
| M13_reverse | CAGGAAACAGCTATGAC |

**C) Nucleotide sequences of primers and probes used in RT-qPCR analysis.**

| **Name** | **Nucleotide sequence (5’ – 3’)** |
| --- | --- |
| **Primers used in RT-qPCR analysis** | |
| **Target gene: *A4GALT*** | |
| A4gt_f | TCCAGGATCGCACTC |
| A4gt_r | GTTGAGGACGTAGCG |
| A4gt_probe | TTCATTGTTCTCAAGAACCTG |
| **Target gene: *B4GALT5*** | |
| B4gt5_f | GGGGACACAGGAAAATACAAG |
| B4gt5_r | CCTCAACAACCTGAACTACTTT |
| B4gt5_probe | CCTCACCACCATCGAGGAGAAGTCCA |
| **Target gene: *GAPDH*** | |
| Gapdh_f | TGGAAAGCTTGTCATCAAC |
| Gapdh_r | GAAGACGCCAGTAGATTCC |
| Gapdh_probe | AGGCCATCACCATCTTCCAG |

**Table SII. RT-qPCR conditions used for quantitative analysis of *A4GALT* and *B4GALT5* transcripts.**

| qPCR system | Reaction format | Reaction volume | Thermal cycling conditions | | | |
| --- | --- | --- | --- | --- | --- | --- |
|  |  | | Parameter | Initial denaturations | PCR (40 cycles) | |
|  |  |  |  |  | Denaturation | Annealing/Extension |
|  |  |  | Temperature [°C] | 95 °C | 95 °C | 60/62* °C |
| 7500 Fast | 96-well plate | 20 µl | Time (mm:ss) | 10:00 | 0:15 | 1:00 |

*****Annealing/extension temperature depended on gene type analysis (60°C for human *A4GALT* or 62°C for hamster *B4GALT5*).
